# Effects of Tasks on Functional Brain Connectivity Derived from Inter-Individual Correlations: Insights from Regional Homogeneity of Functional MRI Data

**DOI:** 10.1101/2024.06.02.597063

**Authors:** Xin Di, Pratik Jain, Bharat B. Biswal

## Abstract

Research on brain functional connectivity often relies on intra-individual moment-to-moment correlations of functional activity, typically using functional MRI (fMRI). Inter-individual correlations are also employed on data from fMRI and positron emission tomography (PET). Many studies have not specified tasks during scanning, keeping participants in an implicit “resting” condition. This lack of task specificity raises questions about how different tasks impact inter-individual correlation estimates. In our analysis of fMRI data from 100 unrelated participants, scanned during seven tasks and in a resting state, we calculated Regional Homogeneity (ReHo) for each task as a regional measure of brain functions. We found that changes in ReHo due to tasks were relatively small compared with its variations across brain regions. Cross-region variations of ReHo were highly correlated among tasks. Similarly, whole-brain inter-individual correlation patterns were remarkably consistent across the tasks, showing correlations greater than 0.78. Changes in inter-individual correlations between tasks were primarily driven by connectivity in the visual, somatomotor, default mode network, and the interactions between them. This subtle yet statistically significant differences in functional connectivity may be linked to specific brain regions associated with the studied tasks. Future studies should consider task design when exploring inter-individual connectivity in specific brain systems.

**Impact Statement:** Inter-individual correlation is increasingly used to estimate brain connectivity, complementing intra-individual correlations in fMRI, particularly for measures like cerebral blood flow obtained via fMRI and PET. However, how task performance affects inter-individual correlations is largely unknown. This study used regional homogeneity as a summary measure of brain functions from task-based fMRI data across eight tasks. The inter-individual correlations were highly similar across tasks, indicating the underlying brain network structure can be inferred under various conditions. Subtle but statistically significant differences in connectivity estimates suggest the functional significance of this approach.

## 1. Introduction

An important goal of brain research is to understand the brain connectivity between remote brain regions (Friston, 2011, 1994). Functional connectivity is mainly studied based on moment-to-moment correlations of brain measures from functional MRI (fMRI) (Biswal et al., 1995, 2010; Fox et al., 2005). An alternative approach is to estimate functional connectivity based on inter-individual correlations of brain functional measures, such as glucose metabolic activity measured using [^18^F]fluorodeoxyglucose ([^18^F]FDG) positron emission tomography (PET) (Horwitz et al., 1984; Metter et al., 1984). Inter-individual covariance of functional structural brain measures have been used for characterizing brain connectivity in both fMRI (Taylor et al., 2012) and PET studies (Di et al., 2017a; Di and Biswal, and Alzheimer’s Disease Neu, 2012).

Previous studies have shown that the inter-individual correlation approach can produce connectivity that are similar to those derived from moment-to-moment correlations using fMRI (Di et al., 2017a) and anatomical connectivity derived from diffusion weighted imaging (Lizarraga et al., 2023). More importantly, the inter-individual approach can take advantage of molecular imaging to measure different aspects of physiology related to neuronal activity (Sala et al., 2023). For example, molecular imaging using PET can measure cerebral blood flow with [^15^O]H_2_O (Frackowiak et al., 1980), glucose metabolism using (F-18)2-fluoro-2-deoxy-D-glucose (FDG) (Phelps et al., 1979), and others (Fang et al., 2021). For fMRI, different types of parameters can also be exploit to reveal different aspects of functions (Taylor et al., 2012). Therefore, although limited by the amount of data, the inter-individual approach could provide unique information regarding brain functional organizations.

Most previous studies do not ask the participants to perform a specific task or place the participants with an explicit “resting” condition. It remains unclear how different task conditions may affect the estimates of functional covariance measures of functional connectivity. Studies using PET have shown that regional functional measures of cerebral blood flow and glucose metabolism were modulated by task conditions in a regionally selective manner. That is, there are certain sets of regions that involved in a particular task performance showed differences in the tasks. These task-specific regional signals may enhance or at least modulate inter-individual correlations between the given region and other brain regions. Alternatively, the enhanced activity might reduce the inter-individual variability, therefore reducing connectivity estimates. In any circumstance, task may lead to changes in inter-individual correlations. However, studies on within individual measures of functional connectivity generally showed that functional connectivity was largely similar across tasks (Cole et al., 2014; Di et al., 2020), and changes in connectivity between task conditions are rather small compared with their absolute values (Cole et al., 2014; Di and Biswal, 2019). The question becomes to what extend is inter-individual connectivity modulated by different task conditions.

Collecting PET data for multiple tasks from the same individuals presents challenges, while obtaining fMRI data, especially when large-scale datasets like the Human Connectome Project (HCP) (Barch et al., 2013) are available, is relatively straightforward. In the HCP dataset, for instance, participants underwent MRI scanning while performing various tasks. We measured functional activations during different tasks using the HCP dataset by using regional homogeneity (ReHo) (Zang et al., 2004), which summarizes brain activations over a period, usually a few minutes. ReHo has demonstrated correlations \with PET and fMRI measures of cerebral blood flow and glucose metabolism (Adhikari et al., 2022; Aiello et al., 2015; Li et al., 2012; Wang et al., 2021). It has also been utilized to derive resting-state functional brain networks using resting-state data (Taylor et al., 2012). In the current study, we further ask how functional connectivity derived from cross-subject correlations of ReHo was modulated by different task conditions.

We first hypothesize that ReHo is a valid measure of brain functions, therefore could show regional alterations in different task conditions according to the task demands. We predict that alteration in ReHo in a task may involve regions that associated with certain task components. We are particularly interested in the extent of changes in ReHo due to task conditions, compared with the variations across different brain regions. Next, we hypothesize that inter-individual correlations of ReHo would be similar across different task conditions. However, we are still interested in specific connections that showed variable connectivity in different task conditions.

## 2. Methods

### 2.1. MRI Data and Task Designs

We conducted an analysis of fMRI data from 100 unrelated participants from the Human Connectome Project (HCP) (Barch et al., 2013), encompassing resting-state scans and seven tasks. Among these participants, 54 were females. Their ages were provided in ranges rather than specific years: 22-25 years (17 participants), 26-30 years (40 participants), 31-35 years (42 participants), and older than 36 years (1 participant).

The participants engaged in seven tasks and a resting state during MRI scanning. These tasks were structured as block designs, each comprising one or more task conditions, corresponding control conditions, and a fixation condition. The resting state is a special task, where participants were instructed to keep their eyes open, avoid falling asleep, and think of nothing in particular. They viewed a “relaxed” fixation, represented by a white cross on a dark background (Smith et al., 2013). With the exception of the language task, which involved auditory stimuli, all tasks utilized visual stimuli or instructions. In each task, participants were required to respond after receiving stimuli, while no response was needed during the resting-state condition. Table 1 provides a breakdown of task parameters for each task.

**Table 1.**
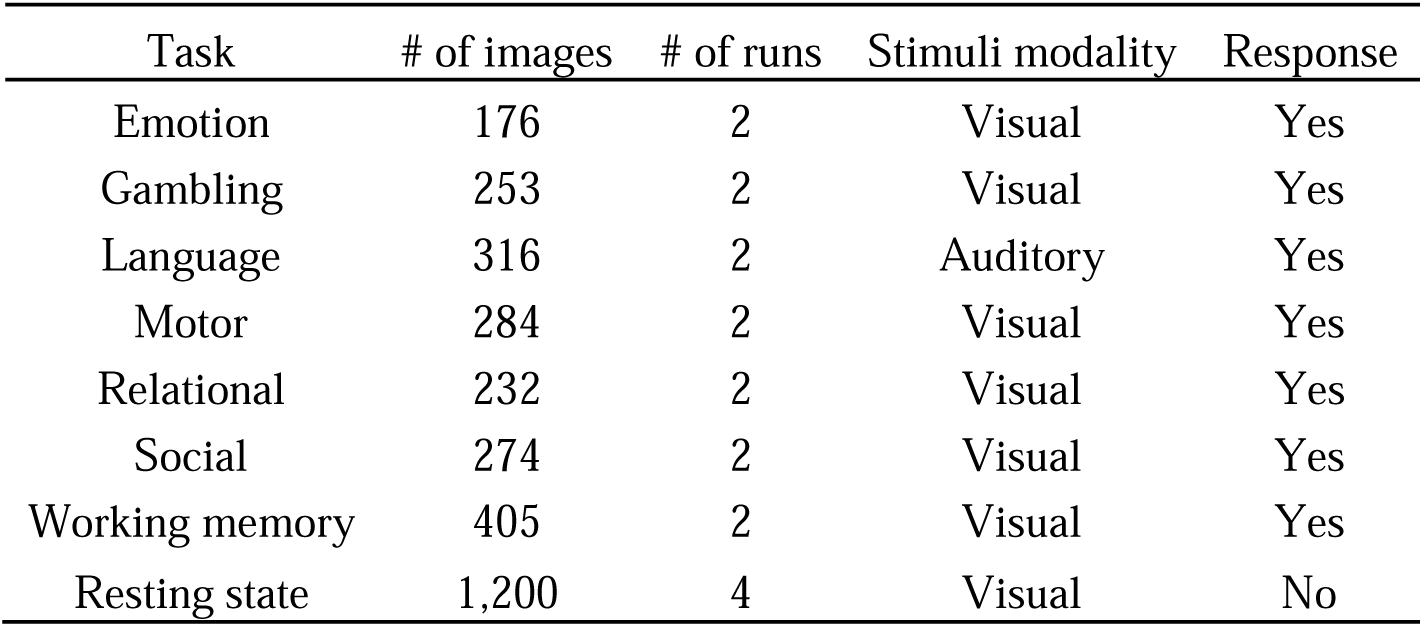
Imaging and task design information for different task conditions.

| Task | # of images | # of runs | Stimuli modality | Response |
| --- | --- | --- | --- | --- |
| Emotion | 176 | 2 | Visual | Yes |
| Gambling | 253 | 2 | Visual | Yes |
| Language | 316 | 2 | Auditory | Yes |
| Motor | 284 | 2 | Visual | Yes |
| Relational | 232 | 2 | Visual | Yes |
| Social | 274 | 2 | Visual | Yes |
| Working memory | 405 | 2 | Visual | Yes |
| Resting state | 1,200 | 4 | Visual | No |

We used the minimally preprocessed data in the current analysis. The MRI data were acquired using Siemens’ standard 32-channel head coil. Each task underwent two fMRI runs with variable durations. One run for each task used right-to-left phase encoding (RL), while the other used left-to-right phase encoding (LR). The resting-state task was scanned in four runs, with two runs for each phase encoding direction. The fMRI scanning parameters were as follows: repetition time (TR) = 720 ms; echo time (TE) = 33.1 ms; flip angle (FA) = 52°; field of view (FOV) = 208 × 180 mm²; slice number = 72; voxel size = 2.0 mm isotropic; multiband factor = 8. The T1-weighted MRI scanning parameters were: TR = 2400 ms; TE = 2.14 ms; FA = 8°; FOV = 224 × 224; voxel size = 0.7 isotropic.

### 2.2. Data Processing and Statistical Analysis

The image processing and data analysis were performed in MATLAB (https://www.mathworks.com), with in-house codes and codes from Statistical Parametric Mapping (SPM12; https://www.fil.ion.ucl.ac.uk/spm/) and Resting-State fMRI Data Analysis Toolkit v1.8 (REST) (Song et al., 2011). BrainNetViewer was used for visualization of brain regions and connections (Xia et al., 2013).

#### 2.2.1. Calculation of ReHo

ReHo was computed as a regional functional measure for each task run (Taylor et al., 2012; Zang et al., 2004). To ensure reliability in correlational analyses, ReHo values were computed across the entire task run without distinguishing between specific experimental conditions within the task (Zuo et al., 2013). Consequently, interpretations should focus on the overall task performed in each run (Table 1) rather than on detailed experimental or control conditions. The whole ReHo calculation process is illustrated in Supplementary Figure S1. Specifically, the preprocessed fMRI images for each run were fed into the ReHo function in REST toolbox to calculate a ReHo map. An intracranial mask was created by combining the intracranial mask from SPM with available functional images for each participant.

Furthermore, the mean ReHo value across all voxels within the intracranial mask was calculated as a global ReHo value, which would be used to standardize regional ReHo values later. Supplementary Figure S2 depicts the averaged ReHo maps for the eight task conditions.

#### 2.2.2. Regions of Interest (ROIs)

We adopted Shaefer’s 100-region parcellation for the cortex (Schaefer et al., 2018). In addition, subcortical regions were also included based on Automated Anatomical Labeling (AAL) atlas (Tzourio-Mazoyer et al., 2002). The 14 included subcortical nuclei were bilateral hippocampus, parahippocampus, amygdala, caudate, putamen, pallidum, and thalamus. For the Schaefer atlas, the regions were separated by the left (#1 to #50) and right (#51 to #100) hemispheres and were further divided into seven functional networks, including visual, sensorimotor, dorsal attention, salience/ventral attention, limbic, control, and default mode networks. In total, 114 regions of interest (ROIs) were used. For each task run, averaged ReHo values were extracted for each ROI, and were divided by the global ReHo value of the participant.

#### 2.2.3. Regional Analysis

Both voxel-wise and ROI-wise analyses were carried out in SPM12 and MATLAB using a repeated-measure generalized linear model (GLM) to assess task-related changes in ReHo (Henson and Penny, 2007). For each of the 100 participants, we concatenated either the mean ReHo images (voxel□wise) or the mean ReHo values within each ROI across seven task conditions (two runs each) and one resting□state condition (four runs), yielding a single dependent□variable vector of length 1,800 [(7 × 2 + 1 × 4) × 100]. The design matrix included eight regressors for the experimental conditions, 100 regressors for subject-specific intercepts, and one regressor coding acquisition phase (LR vs. RL).

First, we specified an omnibus F-contrast to test for any pairwise differences among the eight conditions. Next, we introduced F-contrasts comparing each task condition directly against the resting-state. Finally, we examined the main effect of acquisition phase. Because this phase regressor reached significance (Supplementary Figure S3), we then (1) averaged the LR and RL runs to obtain phase-collapsed estimates and (2) analyzed LR and RL separately to confirm that acquisition order did not confound the observed task effects.

To compare spatial ReHo patterns across task conditions, we first averaged ReHo within each of the 114 ROIs for each task or resting condition by pooling data from all participants and their respective runs (two runs per task, four runs for resting state), yielding an 8 × 114 matrix (eight conditions × 114 ROIs). We then computed Pearson’s correlation coefficients between these condition□specific ROI profiles to produce an 8 × 8 similarity matrix. The same procedure was applied separately to the LR and RL acquisition runs, resulting in one 8 × 8 correlation matrix for each phase.

#### 2.2.4. Inter-Individual Correlation Analysis

Inter-individual correlation matrices among the 114 ROIs were computed using Pearson’s r for each run, yielding a 114 × 114 matrix per run. For task conditions (two runs) and rest (four runs), we first averaged each participant’s run-specific matrices across runs, then averaged those matrices across all 100 participants to produce one 114 × 114 matrix per condition. To quantify similarity between conditions, we extracted the lower-triangular elements of each averaged matrix, vectorized them, and calculated Pearson’s correlations between every pair of condition-specific vectors. In addition, this entire procedure was carried out separately for the LR and RL acquisition phases.

To compare the differences in inter-individual correlations between different task conditions, we adopted an inter-individual psychophysiological interaction (PPI) analysis, which is similar to the regular within participant PPI analysis (Di et al., 2020). The model was constructed on each pair of ROIs, with one as the independent variable and the other as the dependent variable. For both regions, the mean ReHo values across tasks, runs, and participants were concatenated into 1,800 vectors. Before concatenation, the ReHo values were z transformed for each run to normalize the variances. The model included eight regressors representing the task conditions, eight regressors representing ReHo variations of a region in the eight task conditions, 100 regressors represent participant specific effects, and one regressor corresponding to acquisition phase. Please note that this PPI model is an extension to the repeated measure ANOVA model by adding the eight regressors of ReHo values in different task conditions. The regression coefficients of these regressors represent inter-individual correlations between the two regions in a particular condition. Similarly, we defined an F contrast to examine the differences in correlations between any of the two conditions. Further, we also defined F contrasts between each of the task conditions compared with the resting-state condition. Lastly, we performed similar analysis separately for LR and RL runs.

## 3. Results

### 3.1. Regional Effects of ReHo

Whole-brain ReHo maps revealed similar spatial distributions across task conditions (Supplementary Figure S2). As shown in Figure 1A, we present the mean ReHo values across 114 ROI for each run and task conditions. Regionally, posterior regions-including the occipital and parietal regions-generally exhibited higher ReHo values, while frontal, limbic, and subcortical regions showed lower Reho values.

**Figure 1.**
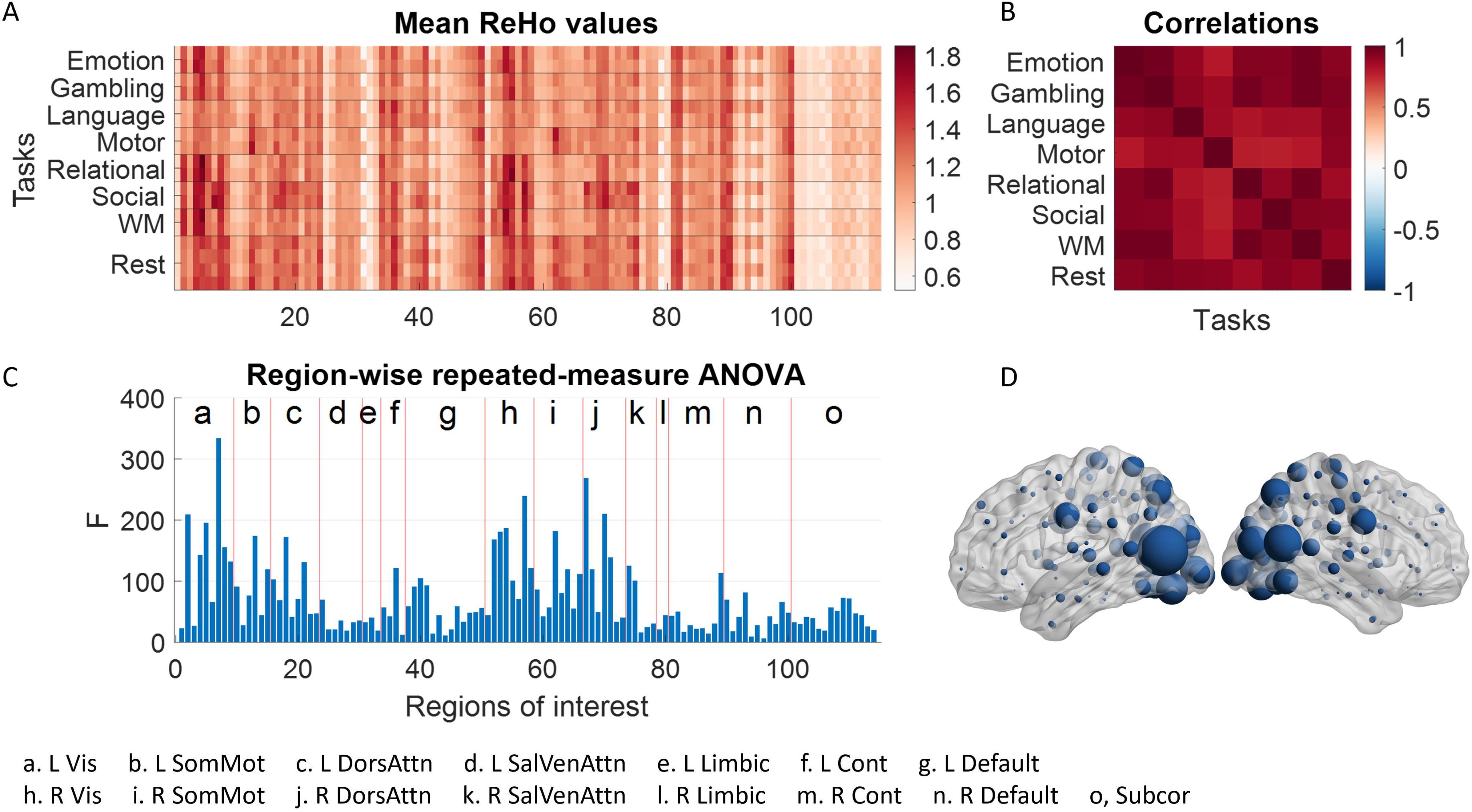
A, Mean regional homogeneity (ReHo) for different brain regions of interest and runs under different task conditions. B, correlations of the mean ReHo values across regions between different task conditions. C, F-statistics of a repeated-measure one-way analysis of variance (ANOVA) with task conditions as the within-subject factor for each of the regions. Vertical lines in C separate different functional networks shown in the bottom. D, Brain representation of the F values in region-wise repeated-measure ANOVA.

Across tasks, the ReHo pattern remained consistent. In addition, there were recurring patterns across the rows, reflecting the effects of acquisition phase (LR vs. RL). To quantify the contribution of these factors, we applied a three-way ANOVA to the matrix shown in Figure 1A. The regional factor accounted for the largest variance (*0.7047*), whereas task conditions and acquisition phase accounted for substantially less variance (0.0019 and 0.022, respectively). Moreover, the mean ReHo patterns across regions were strongly correlated across different task conditions (Figure 1B), with correlation coefficients ranging from *0.7687* to *0.9740*. The lower correlations were primarily observed between the motor task and other task conditions. Similar correlation patterns were found when considering only LR or RL runs separately (Supplementary Figure S4A and S4C).

We utilized repeated measure one-way ANOVA for each region, with task conditions as the within-subject factor and acquisition phase as covariance, to pinpoint regions influenced by task conditions. Figure 1C illustrates the F values for the repeated-measure factor per region, indicating the extent of differences in ReHo among the eight task conditions. Notably, most of the F statistics were statistically significant. However, what’s particularly intriguing are the regions with relatively high F values. Figure 1D reveals that these high F value regions were predominantly situated in the posterior visual regions. Additionally, some sensorimotor network and dorsal attention network regions also exhibited elevated F values compared to other brain regions. Lastly, similar spatial patterns were observed when analyses were conducted separately for LR and RL runs (Supplementary Figure S4B and S4D).

The resting-state condition naturally serves as a control condition for the other tasks. Consequently, we compared each task condition to the resting-state condition (refer to Figure 2). The regions displaying decreased ReHo were primarily within the sensorimotor and dorsal attention networks, notably not the default mode network. Regions exhibiting increased ReHo compared to the resting state varied depending on the tasks. Tasks involving visual stimuli typically led to increased ReHo in visual regions. Additionally, ReHo in certain subcortical regions increased during the gambling task, language task, and notably, the motor task.

**Figure 2.**
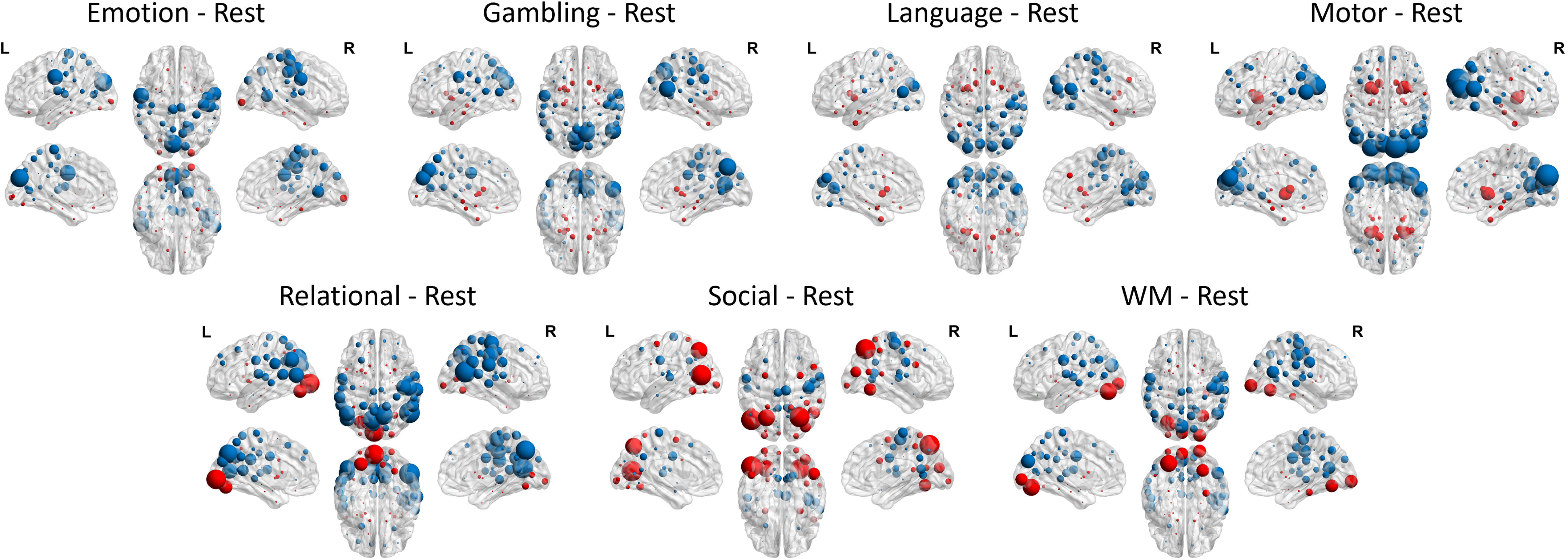
Differences in regional homogeneity (ReHo) between a task condition and the resting-state condition. Red indicates task greater than rest, and blue indicate rest greater than task. The size of the nodes represents F-statistics of the specific contrast without thresholding.

### 3.2. Inter-Individual Correlations

The inter-individual correlation matrices for different task and resting-state runs are depicted in Figure 3. It is evident that these matrices for different task conditions are largely similar. Specifically, within-network connectivity tended to be stronger than between-network connectivity, as seen by the higher values along the diagonal squares. Moreover, there was notable correlation between corresponding networks in the left and right hemispheres. Subcortical regions also displayed strong correlations among themselves, visible in the bottom right corners of the matrices.

**Figure 3.**
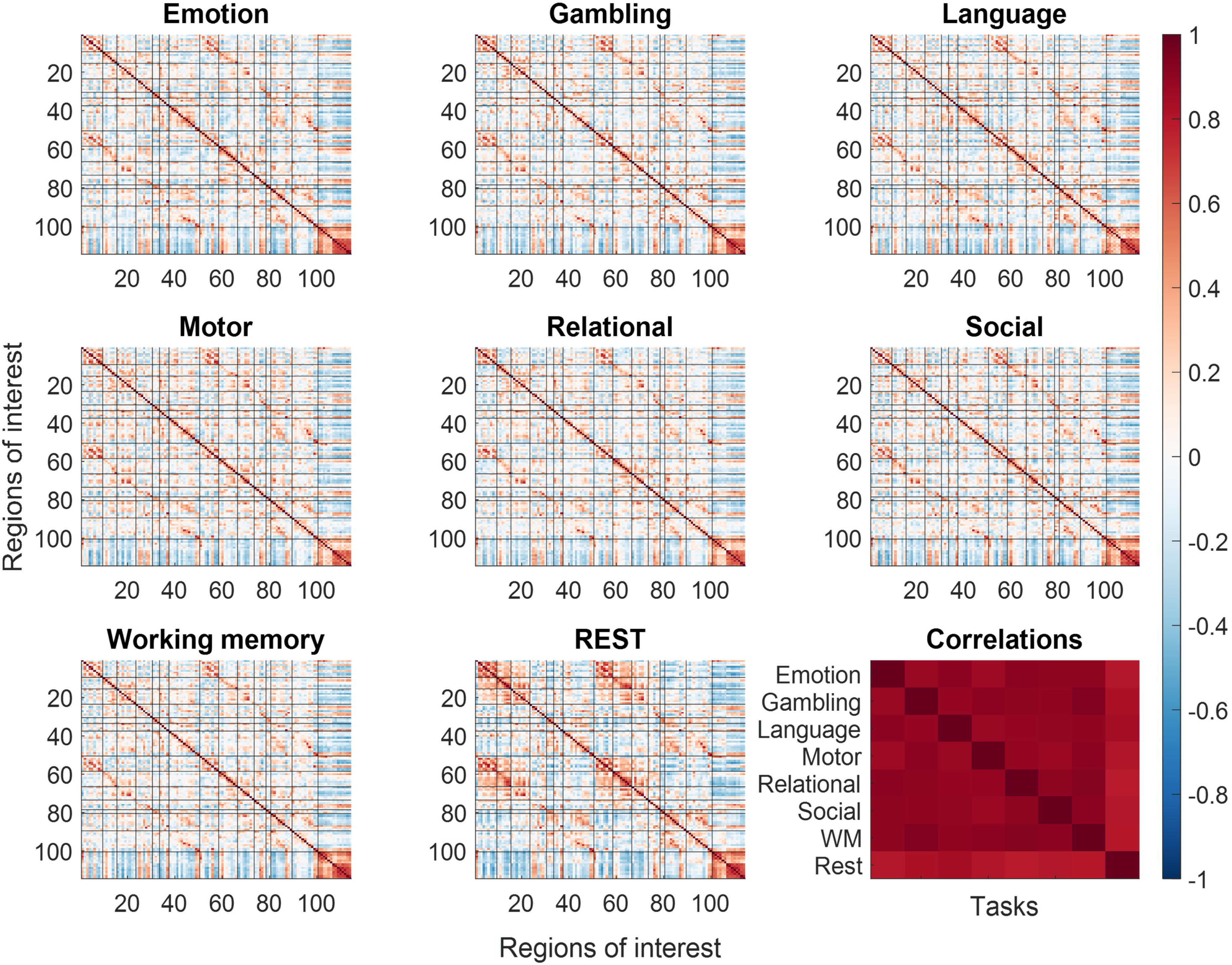
Inter-individual correlation matrices among 114 regions of interest for different task/resting-state conditions. Black lines separate different functional networks. The right most panel in the bottom row shows the correlations of the lower diagonal of the inter-individual correlation matrices among the task and resting-state conditions.

To quantify the similarity across task conditions, we computed correlation coefficients between vectorized upper triangular of the matrices for each task. These matrices exhibited high correlations (ranging from *0.7807* to *0.9386*). Notably, all seven task conditions showed extremely high correlations among each other (around *0.9*), while the correlation between the resting condition and task conditions was slightly lower (around *0.8*). Similar spatial patterns were observed when analyses were conducted separately for LR and RL runs (Supplementary Figure S5).

To pinpoint region pairs whose connectivity is influenced by task conditions, we conducted an inter-subject PPI analysis. This involved employing a repeated-measures ANOVA to detect connectivity that showed significant differences across various task conditions. Although the statistical results differed when one region was treated as the dependent variable and the other as the independent variable, they were generally similar. Consequently, we averaged the lower and upper diagonals to create a symmetrical matrix (refer to Figure 4). The critical F value for p < 0.001 was recorded as 3.50. However, no threshold was applied to the matrix to display all task-modulated connectivity effects. The connections displaying modulated connectivity due to tasks were predominantly within the visual network, somatomotor network, the control network, and among these three networks. Similar spatial patterns were observed when analyses were conducted separately for LR and RL runs (Supplementary Figure S6).

**Figure 4.**
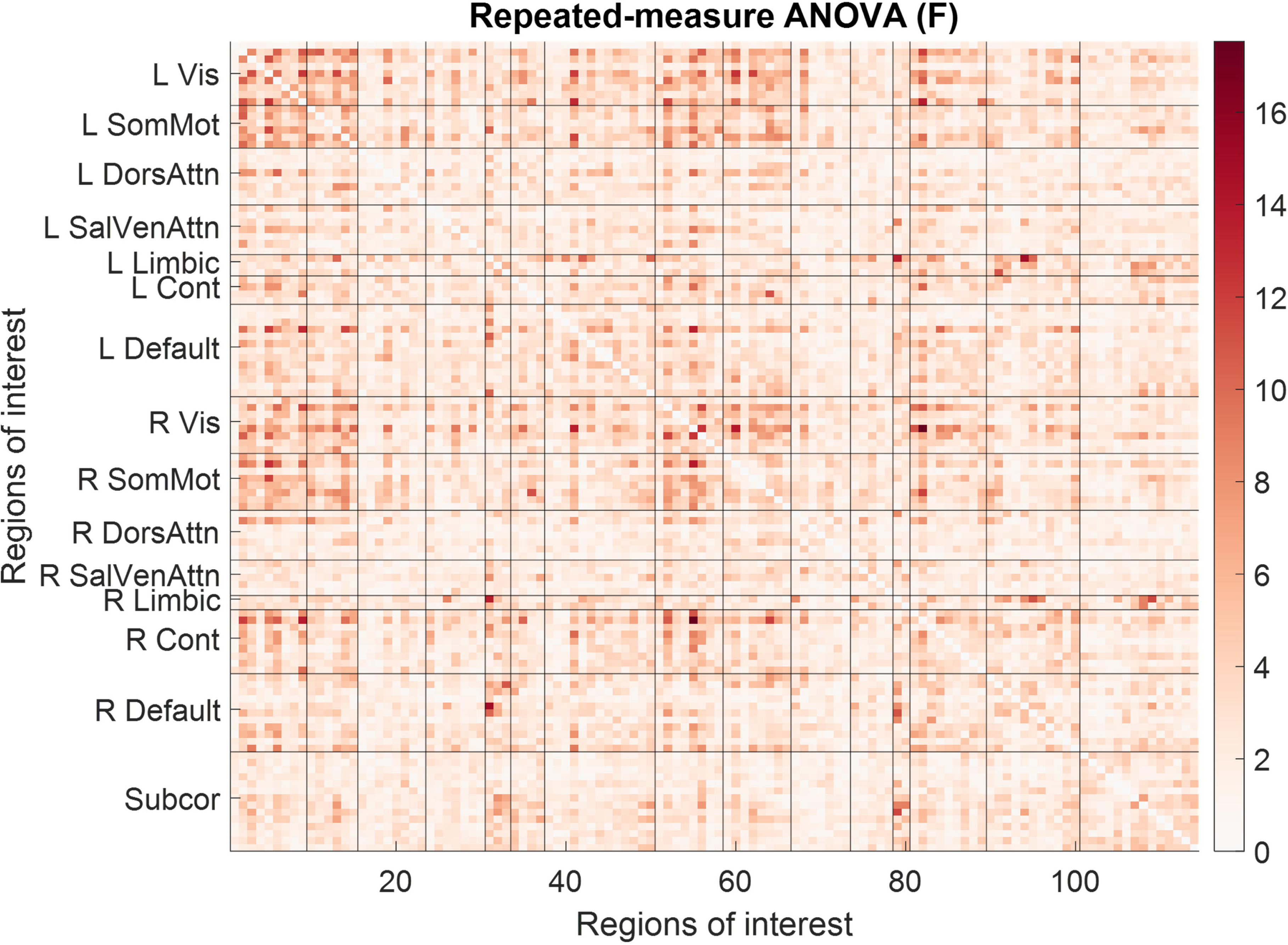
F-statistics representing the changes of inter-individual correlations between any pair of tasks (repeated-measure analysis of variance, ANOVA). Black lines separate different functional networks.

We further contrasted the correlation matrices of each task condition with respect to the resting-state condition (Figure 5). It shows that the general pattern of altered covariance in different task conditions with respect to the resting-state were very similar across tasks. In general, reduced correlations were found within the visual network, sensorimotor network, and between the visual and sensorimotor networks. In contrast, increased correlations were found between the control network and the visual or sensorimotor network, and between the sensorimotor and subcortical networks. For a specific task, there may be unique connectivity modulations, but it is beyond the scope of the current report. To reveal the spatial location of the changes in connectivity for the different tasks, we used BrainNetViewer to show their spatial locations in brain (Figure 6).

**Figure 5.**
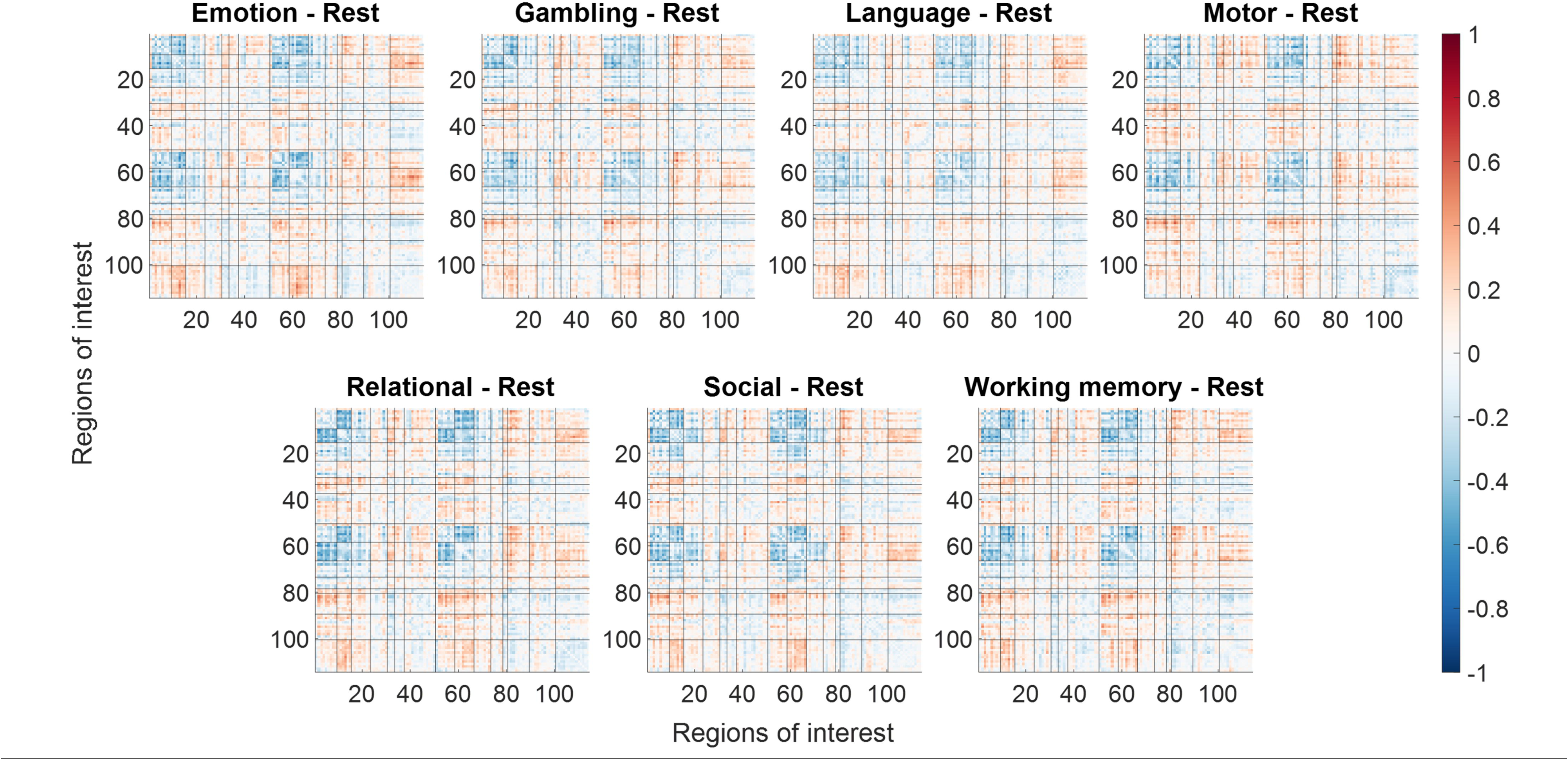
Differences in inter-individual correlations between each task condition with reference to the resting state. Black lines separate different functional networks.

**Figure 6.**
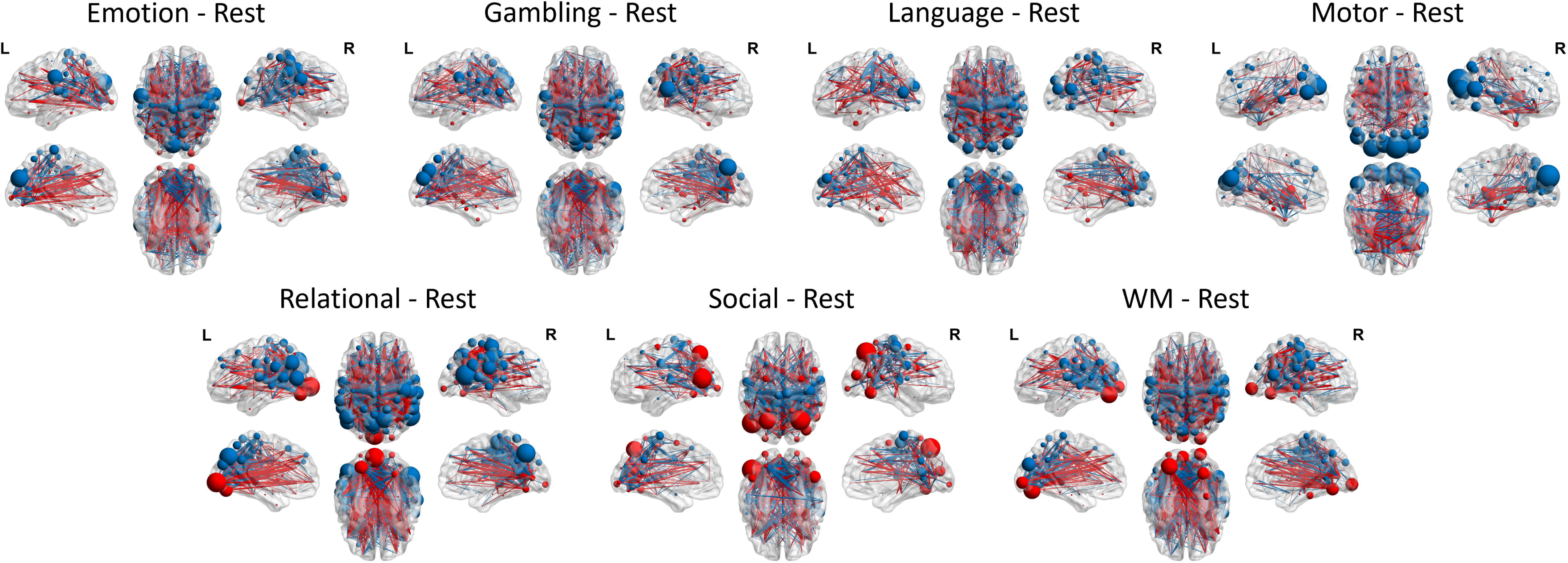
Increased (red) and decreased (blue) inter-individual correlations of regional homogeneity (ReHo) between each task condition compared with the resting condition. The connections were thresholed at F > 10, which correspond to a p value of 0.0016. The size of the nodes represents F-statistics of the specific contrast without thresholding.

## 4. Discussion

By using ReHo as a measure of regional brain functions in a temporal scale of minutes, the current analysis showed to what extent ReHo were modulated by different task conditions and how they may affect the estimates of inter-individual correlation measures of brain functional connectivity. The main results can be summarized as follows: First, the changes in ReHo across different task conditions were relatively small compared with the variations across different brain regions, therefore the spatial correlations of ReHo in different task conditions were highly correlated. Second, inter-individual correlations of ReHo among regions of interest in different task conditions were also highly correlated.

The spatial patterns of ReHo were remarkably consistent among all the task conditions, implying that brain structures likely play a significant role in regulating local synchronizations as assessed by ReHo. Notably, numerous cortical regions exhibited elevated ReHo values, contrasting with the generally lower ReHo observed in subcortical regions. This discrepancy may arise from the greater voxel-level homogeneity within cortical areas compared to the heterogeneous nature of subcortical regions, which encompass various subnuclei. These findings present challenges in utilizing the ReHo method to study subcortical brain activity.

ReHo was developed and mainly used on resting-state fMRI data to reflect functions. Regional ReHo values in resting-state were associated task performance in separate testing (Lee and Hsieh, 2017; Tian et al., 2012), and were also correlated with regional task activations in different tasks (Yuan et al., 2013). But to the best of our knowledge, the current study was the first to demonstrate that ReHo can reflect brain functions in different task conditions. Among a wide range of tasks, we pinpointed brain regions that were more likely to be modulated by the task conditions, which include the visual, sensorimotor, and dorsal attention networks. It makes sense because the unimodal regions directly receive input from outside world and generate response, which varied by different task conditions. In contract, the default mode network regions were among the regions that showed the smallest changes in ReHo values in different task conditions. And by directly contrasting each task condition with the resting-state condition, we further confirmed the regions showed reduced activity in tasks compared with resting-state were mostly sensorimotor, dorsal attention, and visual regions, but not the default mode regions. This result is a little in contrast with PET studies (Shulman et al., 1997), and may reflect the dissociations between glucose metabolism and BOLD oxygenation in the default mode regions (Stiernman et al., 2021).

The regions displaying increased task modulation primarily belong to unimodal brain networks. For instance, the visual networks exhibited increased ReHo values during tasks involving constant visual stimuli, like the emotion task, relational task, social task, and working memory task. Similarly, the auditory region showed elevated ReHo values during the language task, aligning with the auditory nature of the stimuli in this task. Interestingly, the motor regions demonstrated a reduction in ReHo during tasks compared to resting-state conditions. This suggests that task execution involves specific regions within the sensorimotor network, leading to decreased activity in other sensorimotor network regions. Additionally, subcortical regions displayed elevated ReHo values during certain tasks such as the gambling task, language task, motor task, and social task, which is logical given their involvement in motor and reward-related functions. A noteworthy practical implication is that although ReHo values are generally low in subcortical regions, engaging in certain tasks can boost brain activation levels in these regions, thereby enhancing signals within them.

The inter-individual correlations of ReHo were also highly similar across the task conditions, which is consistent with studies of within-individual connectivity (Cole et al., 2014; Di et al., 2020; Krienen et al., 2014). This is also consistent with the high similarity of regional activity, and suggest a common brain network structure may contribute to the highly correlated connectivity structure.

Notable differences in local inter-individual connectivity were also present. The resting state was the most deviate condition and showed relatively lower correlations with all the seven tasks. The connectivity that was modulated by different task conditions can be further categorized into two groups. Firstly, the connectivity within the visual and sensorimotor networks and the connectivity between the two networks typically showed reduced connectivity in any task conditions compared with the resting state. This is in line with within-individual connectivity changes, where similar connectivity reduction are seen when comparing task conditions with the baseline fixation condition within a task run (Di et al., 2020, 2017b) a separate resting-state run (Cole et al., 2014). Second, the connectivity between the control network and regions from the visual, sensorimotor, and subcortical networks showed increased connectivity in various tasks compared with the resting-state. This is also in line with within-individual comparison between tasks and resting-state (Cole et al., 2014), which in line with the role of the control network as a flexible hub that dynamically connected to other brain regions upon task demands (Cole et al., 2013). Again, a notable absence was the connectivity involving the default mode network, which didn’t show much task modulations on connectivity.

Task modulation provides stronger evidence of inter-individual correlations reflecting functional connectivity, contrasting previous mere observations of such correlations. The current findings also carry practical implications for utilizing inter-individual correlation in studying functional connectivity. Specifically, they demonstrate that the overall connectivity pattern remains largely consistent across different task conditions, indicating the feasibility of studying whole-brain functional connectivity in diverse psychological states.

However, subtle yet statistically significant connectivity changes also highlight the importance of considering task design, especially for regions with well-known functions. For instance, if a region of interest is typically activated during a specific task, employing that task may enhance activity and impact correlation estimates with other regions. This consideration is particularly relevant for subcortical regions, where activity is generally weak during resting-state. Additionally, interpretations of results involving sensorimotor regions require attention, as regional activity often decreases during tasks compared to resting-state conditions.

The current analysis was centered on blood-oxygen-level dependent (BOLD) signals, leaving uncertainties about generalizing these results to molecular imaging. Task-induced signal changes typically remain under 5% in a standard 3T MRI scanner (Zandbelt et al., 2008; van der Zwaag et al., 2009), possibly resulting in limited ReHo alterations. Contrastingly, changes in regional cerebral blood flow and glucose metabolism can lead to significant signal fluctuations. For instance, cerebral blood flow changes measured by arterial spin labeling (ASL) MRI have been reported to range from 10% to 26% during a 2-back task (Kim et al., 2006), while glucose metabolic activity changes vary from 18% to 28% across different tasks and studies (Rischka et al., 2018; Vlassenko et al., 2006). Task-related modulation may be more pronounced in PET imaging due to substantial signal changes, necessitating further systematic investigation.

Several methodological considerations merit discussion. First, ReHo was calculated across entire task runs, encompassing transitions between different task blocks. Given that task-induced changes in BOLD signals are typically small compared to intrinsic fluctuations (typically < 5%), the observed decrease in ReHo during tasks compared to resting-state suggests that task transitions did not inflate ReHo values under task conditions. Second, the varying number of time points across tasks is a limitation. Although this is suboptimal, the ReHo measure is not dependent on the number of time points, which ensures valid comparisons across tasks. Arbitrarily matching the number of time points would likely reduce the reliability of tasks with longer durations. Additionally, we reanalyzed the data using only one resting-state run and found results comparable to those derived from all four runs, indicating that differences in the number of time points are unlikely to impact the conclusions of this study. Third, it only relied on ReHo to depict regional activity, while alternative methods like the amplitude of low-frequency fluctuations (ALFF) exist (Zang et al., 2007), which could impact inter-individual connectivity patterns (Taylor et al., 2012). However, exploring these different measures falls beyond the scope of the present analysis.

## 5. Conclusion

ReHo can be utilized with task fMRI data to capture stable brain functions over minutes. Nevertheless, ReHo values exhibit smaller differences across various task and resting-state scenarios compared with cross-region variations. Inter-individual functional connectivity can be inferred from ReHo values, with whole-brain correlation matrices showing consistency across task and resting-state conditions. Notably, connectivity strength changes were primarily observed between the resting state and other task conditions. While the choice of task may not significantly alter the overall connectivity matrix estimate, it can influence specific connectivity patterns with regions involved in task modulation.

## Author Contribution

X.D., Conceptualization, Data curation, Formal Analysis, Visualization, Methodology, Funding acquisition, Writing – original draft.

P.J., Data curation, Formal Analysis, Writing – review & editing.

B.B.B., Conceptualization, Funding acquisition, Writing – review & editing.

## Acknowledgement

The initial idea was developed through discussions with the Molecular Connectivity Working Group (MCWG). The authors would like to thank Dr. Rui Yuan, Marielys Reyes Castillo, and Pushti Shah for their assistance with data curation and image processing. A preprint of this manuscript has been submitted to bioRxiv for scientific communication. The citation details are as follows: Di X, Pratik Jain, Bharat B Biswal (2024). Effects of Tasks on Functional Brain Connectivity Derived from Inter-Individual Correlations: Insights from Regional Homogeneity of Functional MRI Data. bioRxiv 2024.06.02.597063; doi: https://doi.org/10.1101/2024.06.02.597063

