## Supplementary materials for "Effects of Tasks on Functional Brain Connectivity Derived from Inter-Individual Correlations: Insights from Regional Homogeneity of Functional MRI Data"

This document provides supporting information for the manuscript titled “Effects of Tasks on Functional Brain Connectivity Derived from Inter-Individual Correlations: Insights from Regional Homogeneity of Functional MRI Data.” It contains Supplementary Figures S1 through S6.

**Supplementary Figure S1** Flowchart of regions homogeneity (ReHo) calculation.

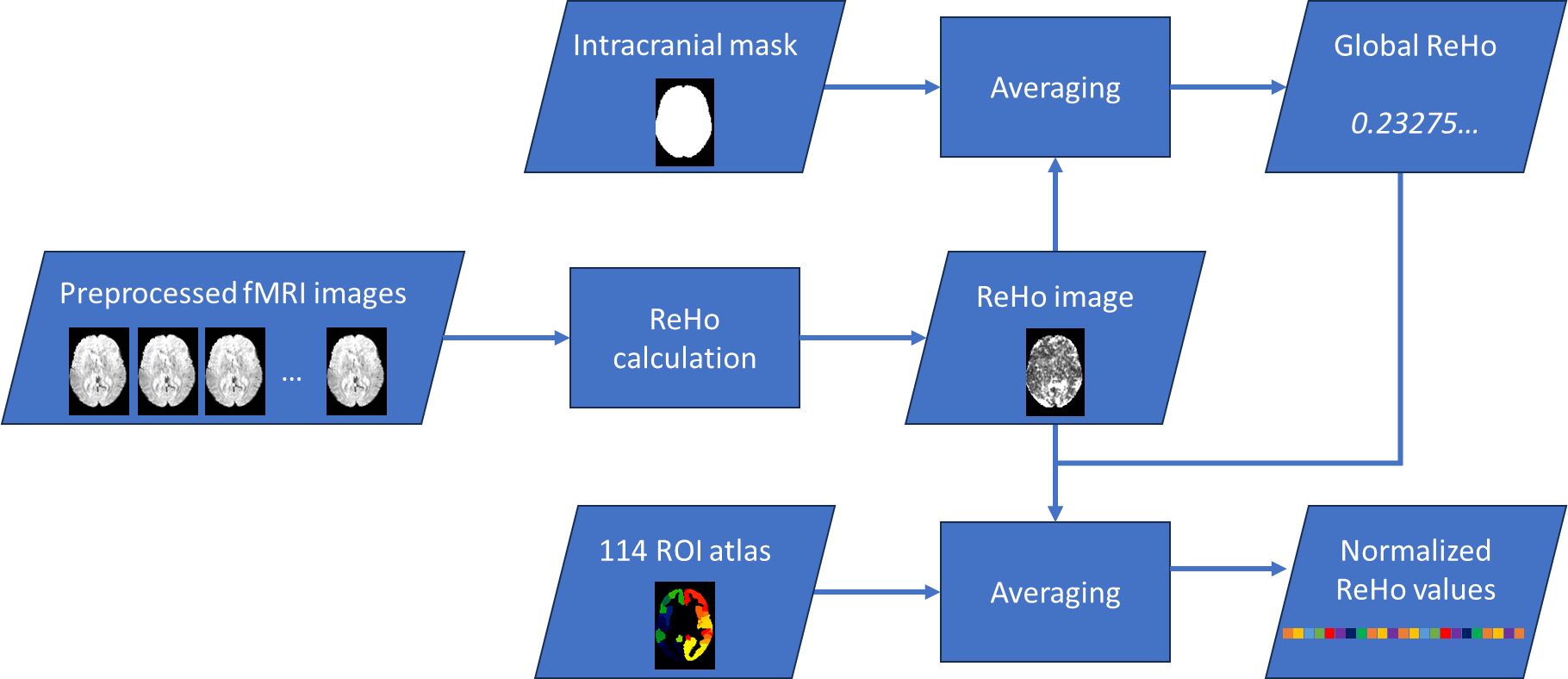

**Supplementary Figure S2** Averaged regional homogeneity (ReHo) images for each task across individuals and runs.

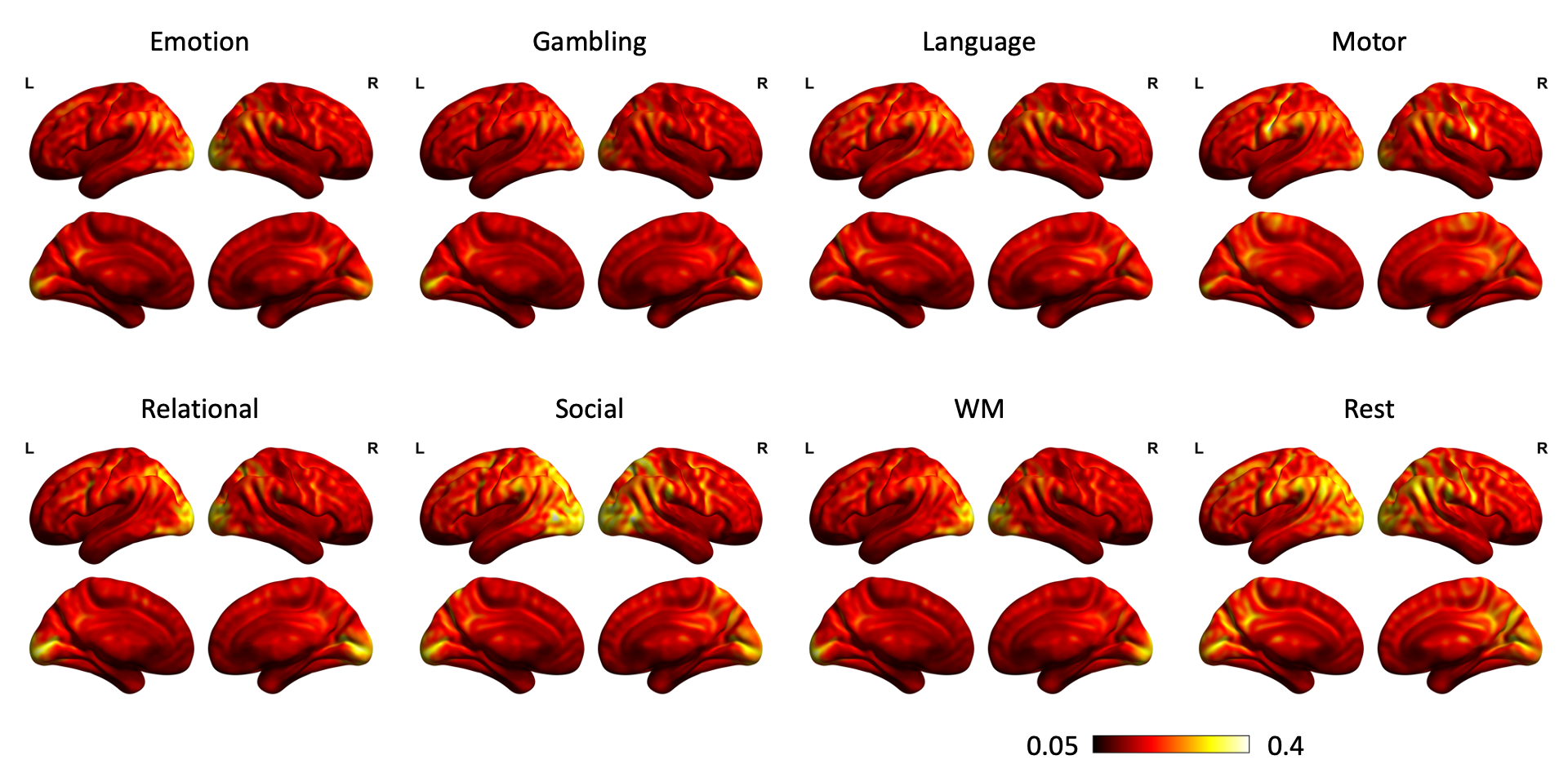

**Supplementary Figure S3** Voxel-wise effects of phase (t) in the repeated-measures ANOVA of ReHo values. Warm colors indicate greater ReHo in the LR phase compared to RL, while cool colors indicate the opposite (RL > LR). The maps were thresholded at |t| > 3, approximately corresponding to an uncorrected p-value of < 0.001.

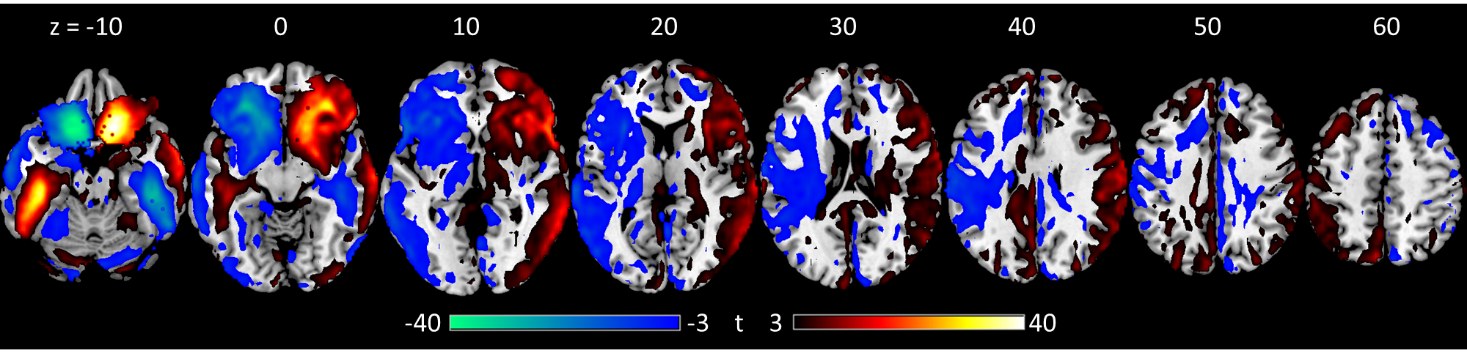

**Supplementary Figure S4** Correlations of the mean ReHo values across regions between different task conditions for LR runs (A) and RL runs (C). F statistics of a repeated-measure one-way analysis of variance (ANOVA) with task conditions as the within-subject factor for each of the regions for the LR runs (B) and RL runs (D). The vertical lines in B and D separate different functional networks shown in the bottom.

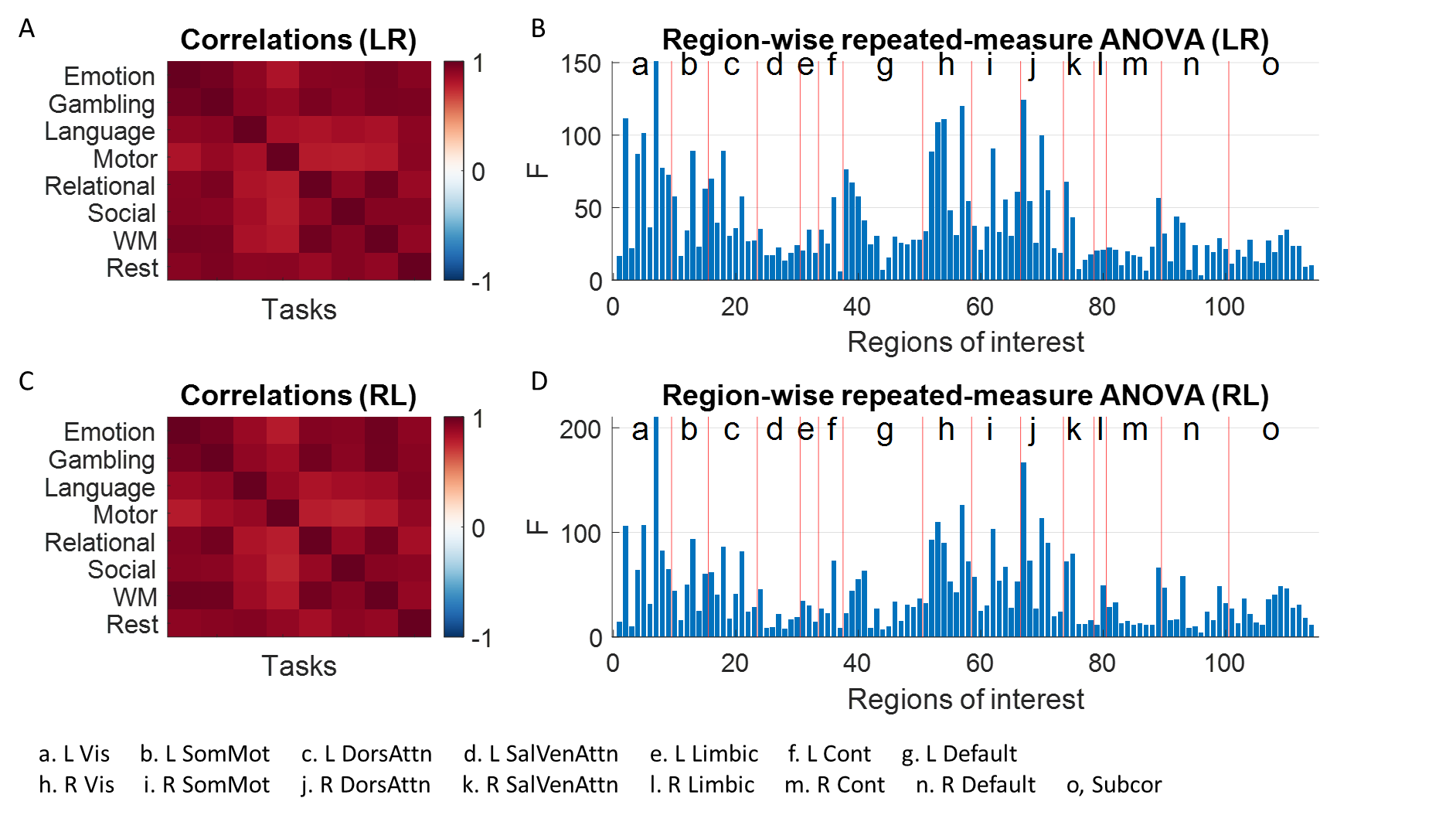

**Supplementary Figure S5** Correlations of the lower diagonal of the inter-individual correlation matrices among the task and resting-state conditions for the LR (left) and RL (right) runs.

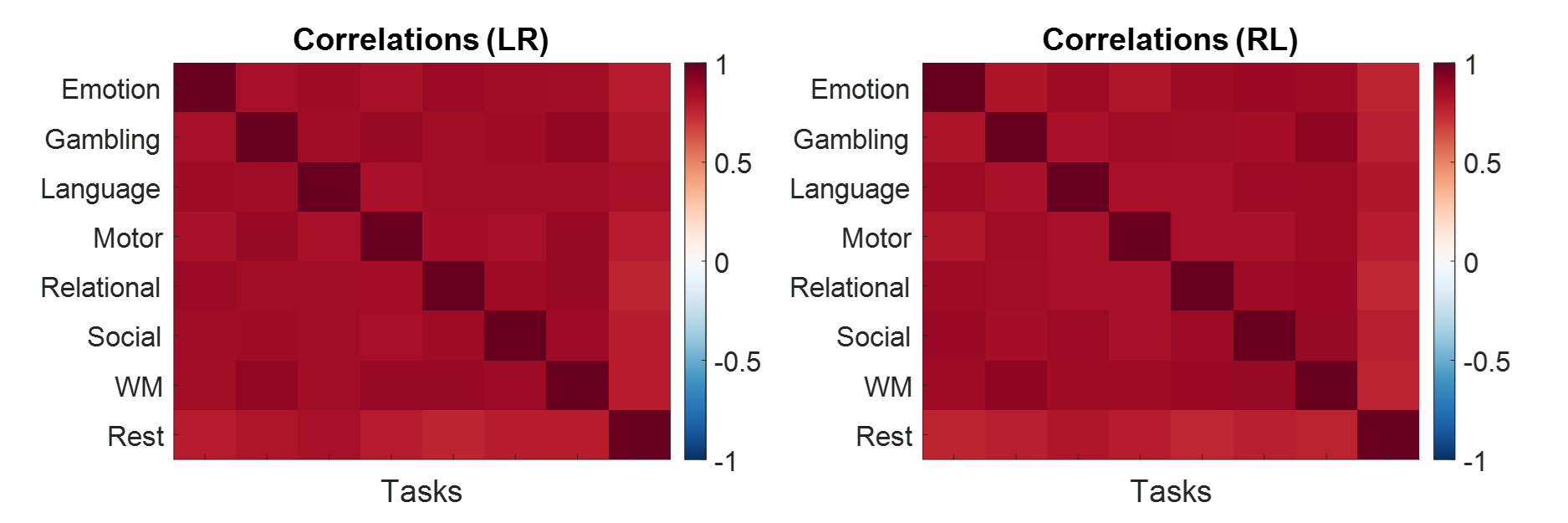

**Supplementary Figure S6** F-statistics representing changes in inter-individua correlations between any pair of tasks (repeated-measure analysis of variance, ANOVA) are shown separately for LR (left) and RL (right) runs. Black lines separate different functional networks.

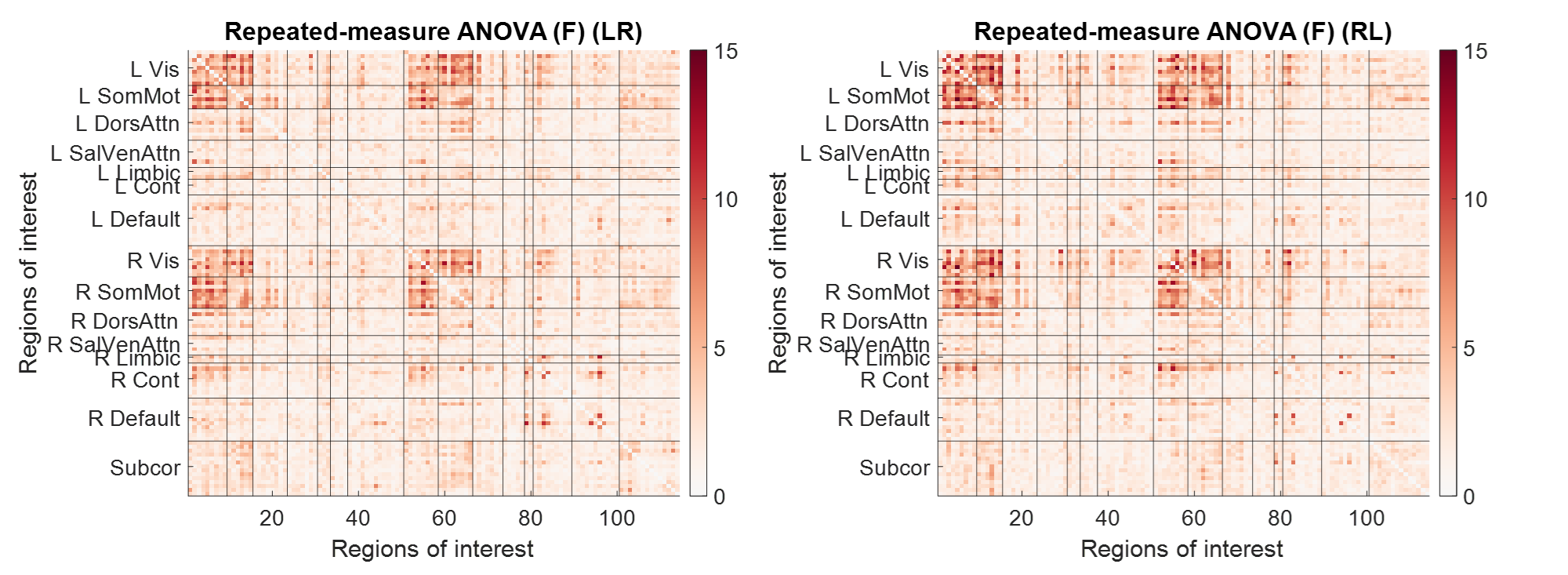
